# Metabolic engineering of *Acinetobacter baylyi* ADP1 for efficient utilization of pentose sugars and production of glutamic acid

**DOI:** 10.1101/2025.03.16.643438

**Authors:** Jin Luo, Elena Efimova, Ville Santala, Suvi Santala

## Abstract

Efficient utilization of pentose sugars is critical for advancing sustainable biomanufacturing using lignocellulosic biomass. However, many potential host strains capable of consuming glucose and lignin-derived monomers are unable to utilize pentose sugars. Here, we engineered *Acinetobacter baylyi* ADP1 for the utilization of D-xylose and L-arabinose. We first modelled different pentose utilization pathways using flux balance analysis to choose the most optimal pathway. A novel marker-free approach facilitated the seamless integration of the pentose catabolic gene clusters of the selected Weimberg pathway into the *A. baylyi* genome, generating strains capable of efficiently utilizing both D-xylose and L-arabinose as sole carbon sources without any additional engineering or adaptation. For D-xylose, the cells achieved the highest growth rate (µ=0.73 h^-1^) reported to date for non-native hosts engineered for pentose utilization. For L-arabinose, a growth rate of µ=0.35 h^-1^ was achieved, which also surpassed the growth rate on a native substrate of *A. baylyi*, glucose (µ=0.31 h^-1^). Importantly, pentose utilization occurred simultaneously with glucose utilization. We then applied metabolic flux analysis using ^13^C labeled xylose to reveal D-xylose metabolism in the engineered strain. To demonstrate the potential for value-added bioproduct synthesis, L-glutamate was selected as a target compound. Deletion of *sucAB* and *gabT*, and the overexpression of *gdhA* enabled the production of L-glutamate. With the engineered strain, a carbon yield of 34% during co-utilization with succinate and 70% via whole-cell catalysis using resting cells was achieved. This study establishes a robust platform for pentose utilization and value-added bioproduct synthesis in *A. baylyi* ADP1 and highlights the potential for further metabolic optimization.

## 1. Introduction

Harnessing the potential of lignocellulosic biomass for bioproduction is crucial for advancing sustainable manufacturing and mitigating the environmental impact of conventional production processes (Long et al., 2024). However, achieving economic viability remains challenging due to the incomplete conversion of lignocellulosic components. Hemicellulose, an abundant component of lignocellulosic biomass, represents a vast and underutilized resources (Geng et al., 2022; Narisetty et al., 2022). This heterogeneous polysaccharide is predominantly composed of pentose sugars, such as D-xylose and L-arabinose, with lesser amounts of hexose sugars, including D-glucose (Scheller and Ulvskov, 2010). Valorization of hemicellulose is essential for maximizing biomass utilization and enhancing the efficiency of bioproduction processes.

Several industrially relevant microorganisms, such as *Corynebacterium glutamicum* and *Saccharomyces cerevisiae*, naturally lack the ability to metabolize pentoses, while others, like *Escherichia coli*, face challenges in efficiently utilizing mixed sugars such as D-glucose and D-xylose due to carbon catabolite repression (Choi et al., 2019; Kaplan et al., 2024; Qiu et al., 2023; Singh et al., 2008). Metabolic engineering has been widely applied to overcome these limitations and enable efficient xylose utilization (Choi et al., 2019; Dvořák et al., 2024; Li et al., 2019; Qiu et al., 2023). However, the direct utilization of cellulosic or hemicellulosic hydrolysates is further hindered by the presence of various inhibitors, including furan derivatives and lignin-derived aromatics, which pose challenges for bioproduction (Guo et al., 2022; Linh et al., 2017; Mills et al., 2009; Narisetty et al., 2022). Selecting a host strain capable of tolerating and utilizing these inhibitors offers a potential solution.

*Acinetobacter baylyi* ADP1 has emerged as a promising host for diverse applications in synthetic biology and industrial biotechnology (Biggs et al., 2020; de Berardinis et al., 2009; Santala and Santala, 2021; Young et al., 2005). This gram-negative, strictly aerobic soil bacterium is known for its exceptional catabolic versatility, allowing it to metabolize a wide range of substrates, and its natural competence and recombination machinery, which facilitates efficient genetic engineering. Notably, *A. baylyi* ADP1 is capable of degrading lignin-derived aromatics, making it an attractive candidate for lignin valorization (Beckham et al., 2016; Salvachúa et al., 2015). It demonstrates high tolerance to aromatic compounds and has been engineered to produce various chemicals from lignin-derived aromatics (Arvay et al., 2021; Luo et al., 2019; Meriläinen et al., 2024; Salmela et al., 2019). Additionally, the bacterium has shown the ability to detoxify inhibitory furan derivatives which are commonly present in lignocellulosic hydrolysates (Liu et al., 2024a, 2024b). These traits make *A. baylyi* ADP1 an excellent platform for hemicellulose valorization by detoxifying/metabolizing the furan derivatives and lignin-derived aromatics from the hydrolysate. However, a critical limitation of *A. baylyi* ADP1 is its inability to metabolize relevant pentose sugars, namely xylose and arabinose.

Xylose metabolism in microorganisms has been well-studied and is summarized in numerous review articles (Kim and Woo, 2018; Kuschmierz et al., 2022; Li et al., 2019; Zhao et al., 2020). It can be broadly classified into two primary strategies based on whether pentose sugars are oxidized to the corresponding sugar acids, with the isomerase and Weimberg pathways serving as key representatives. The isomerase pathway, the most widely applied, converts xylose into xylulose, which subsequently enters the pentose phosphate pathway to generate fructose-6- phosphate and glyceraldehyde-3-phosphate. Xylulose can also be processed into C2 and C3 molecules via xylulose-1/5-phosphate through alternative routes. The Weimberg pathway, in contrast, bypasses phosphorylation steps, converting xylose into 2-keto-3-deoxy-xylonate, which directly enters the tricarboxylic acid (TCA) cycle as α-ketoglutarate or is further cleaved into C2 and C3 molecules via the Dahms pathway. Theoretical product yields are highly dependent on the chosen pathways, making the selection of the appropriate pathway/product critical for optimizing bioprocess efficiency.

In this study, we first used flux balance analysis (FBA) to select the optimal pentose utilization pathway in terms of carbon yield. We then introduced the selected Weimberg pathway into *A. baylyi* ADP1 by integrating the D-xylose and L-arabinose catabolic gene clusters from *A. baumannii* (Alberti et al., 2023) into the *A. baylyi* ADP1 genome. This was achieved using a novel genetic integration method that takes advantage of the bacterium’s natural competence and high recombination activity. We characterized the pentose utilization capacity of the engineered strain and performed carbon flux analysis to better understand the D-xylose metabolism. Given that the Weimberg pathway is advantageous for producing chemicals derived from α-ketoglutarate, we further engineered the strain by blocking the TCA cycle to produce L-glutamate from D-xylose, as a demonstration of its potential to be the platform for hemicellulose valorization.

## 2. Results and discussion

### 2.1. Selection of the optimal pentose utilization pathway

The selection of an efficient pentose utilization pathway is crucial for the economic feasibility of lignocellulose-based biomanufacturing. Multiple pentose utilization pathways have been studied across different microbes (Kim and Woo, 2018; Li et al., 2019) and can be classified based on whether pentose sugars are oxidized to pentose acids (Figure S1): non-oxidative pathways, including the isomerase, xylose reductase-xylitol dehydrogenase (XR-XDH), synthetic xylulose-1-phosphate (X1P), and phosphoketolase pathways, process pentose sugars without oxidation; in contrast, oxidative pathways, such as the Weimberg and Dahms pathways, involve oxidation of pentoses to pentose acids (e.g., xylonate). The steps of each pathway are summarized in Table S1.

Recent studies introduced the isomerase pathway into *Pseudomonas putida* (Dvořák et al., 2024; Elmore et al., 2020a), a bacterium closely related to *A. baylyi* ADP1 in its ability to metabolize lignin-derived aromatics and tolerate inhibitors found in lignocellulosic hydrolysates. However, unlike the isomerase pathway and other non-oxidative pathways that produce C2 or C3 metabolites as the end products, the Weimberg pathway converts D-xylose directly to the TCA cycle intermediate α-ketoglutarate. To compare pathway efficiencies, we performed FBA to calculate the theoretical yields of different TCA cycle intermediates from D-xylose. The results showed that the Weimberg and phosphoketolase pathways outperform other pathways in α-ketoglutarate production, achieving a 100% molar yield (Table S2). α-Ketoglutarate serves as an important precursor for the biosynthesis of a wide range of valuable chemicals (Chae et al., 2017; Lee et al., 2019). Furthermore, the Weimberg pathway bypasses the tightly regulated pentose phosphate and glycolysis pathways, reducing competition for metabolic flux with other cellular processes and enabling more efficient carbon flow toward desired products. Notably, this pathway was recently discovered and characterized in *A. baumannii* (Alberti et al., 2023), a species belonging to the same genus as *A. baylyi* ADP1, suggesting strong compatibility with *A. baylyi* ADP1. These features make the Weimberg pathway attractive for heterologous expression in *A. baylyi* ADP for the production of valuable compounds derived from TCA cycle.

The Weimberg pathway in *A. baumannii* consists of two clusters (Figure 1). Cluster I is responsible for the degradation of L-arabinonate, encoding enzymes such as L-arabinonate dehydratase (AraC1), L-2-keto-3-deoxyarabinonate dehydratase (AraD1), and α-ketoglutaric semialdehyde dehydrogenase (AraE1), catalyzing the stepwise conversion of L-arabinonate to α-ketoglutarate. This cluster also includes a specific uptake transporter (AraI1) and a regulatory protein (AraR1) that acts as an activator of the pathway. Cluster II, on the other hand, is specialized for the degradation of D-xylonate and D-ribonate, encoding homologous enzymes (AraC2, AraD2, and AraE2), with AraE2 functioning similarly to AraE1 in the final oxidation step. This cluster also contains a different transporter (AraI2), and its regulatory protein (AraR2) acts as a repressor.

**Figure 1.**
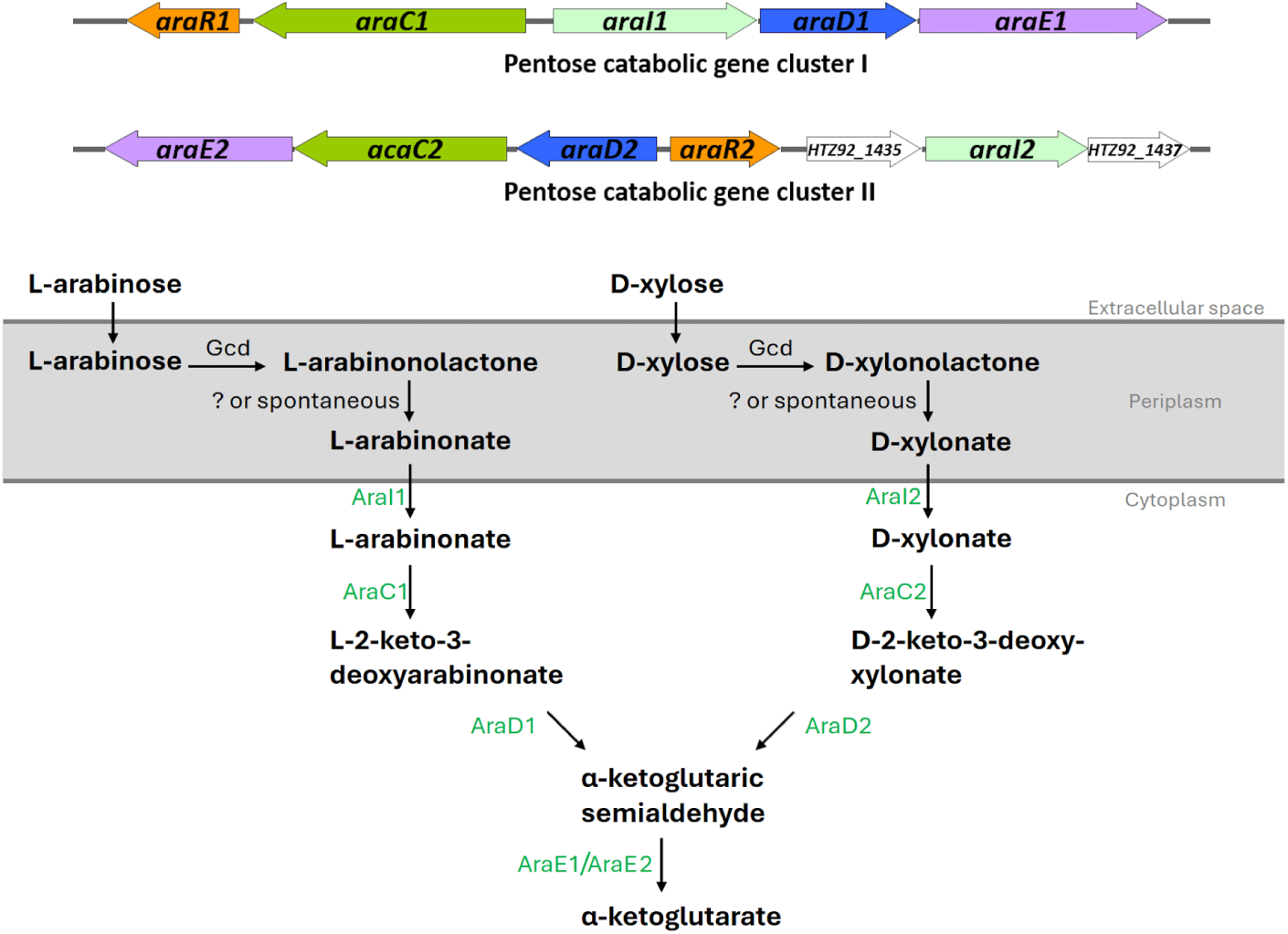
Organization of the pentose catabolic gene clusters in *A. baumannii* (as described in the reference (Alberti et al., 2023)) and the proposed heterologous degradation pathways of D-xylose and L-arabinose in *A. baylyi* ADP1. Abbreviations: Gcd, glucose dehydrogenase; AraI1, L- arabinoate transporter; AraI2, D-xylonate transporter; AraC1, L-arabinonate dehydratase; AraC2, D-xylonate dehydratase; AraD1, L-2-keto-3-deoxyarabinonate dehydratase; AraD2, D-2-keto-3- deoxy-xyolonate dehydratase; AraE1/AraE2, α-ketoglutaric semialdehyde dehydrogenase. The conversion of L-arabinolactone/D-xylonolactone to L-arabinonate/D-xylonate is spontaneous or catalyzed by uncharacterized lactonases.

In *A. baylyi* ADP1 genome, homologs of the aforementioned enzymes can be found, except for AraD1 and AraD2. The conversion of pentose to pentonate is expected to be achieved by the nonspecific activity of the native membrane-bound glucose dehydrogenase Gcd (Santala et al., 2018), followed by spontaneous lactonolysis. Thus, it was hypothesized that the expression of *araD1* and *araD2* could enable the utilization of L-arabinose and D-xylose, respectively. To test this hypothesis, the gene *araD1* was cloned into the plasmid pBAV1C-chn (Luo et al., 2019) under the control of a cyclohexanone-inducible promoter and introduced into wild-type *A. baylyi* ADP1, the resulting strain was designated as ASA546. However, the expression of *araD1* alone did not enable the cells to grow on L-arabinose. The finding suggests that the inability of *A. baylyi* to grow on pentose sugars is not only due to the absence of L-arabinonate/D-xylonate dehydratase (AraD) homologs. Nevertheless, it was observed that both L-arabinose and D-xylose could be oxidized to arabinonate and xylonate, respectively, indicating that the integration of the additional genes in the clusters may achieve pentose utilization.

### 2.2 Integration of the pentose catabolic gene clusters into the genome of *A. baylyi*

The natural competence and high recombination efficiency of *A. baylyi* ADP1 make it an excellent platform for genetic engineering applications (Barbe, 2004; Bedore et al., 2023; Santala and Santala, 2021). Taking advantages of these capabilities, we developed a novel, marker-free methodology to integrate non-native catabolic gene clusters into the genome of *A. baylyi* ADP1 (Figure 2A). This approach involves natural transformation by introducing a heterologous gene cluster flanked by homologous sequences into a culture containing both native and non-native carbon sources. During incubation, transformants capable of utilizing the non-native substrate are enriched, enabling their selection based on metabolic capability. Given that *A. baylyi* and *A. baumannii* belong to the same genus and share compatible genetic elements, integrating entire catabolic gene clusters is advantageous for maintaining balanced enzyme cascades and optimizing metabolic functionality.

**Figure 2.**
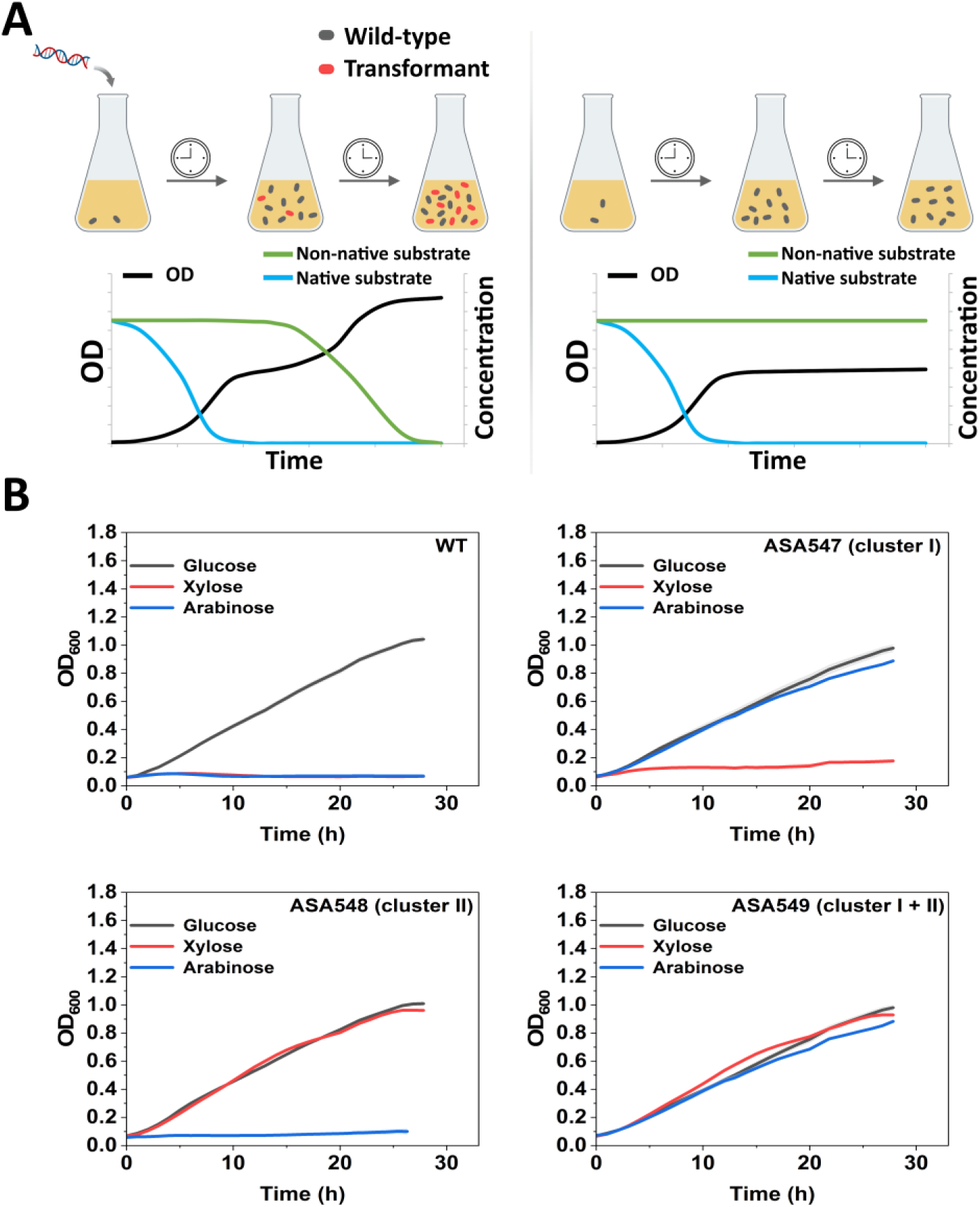
Introduction of the pentose utilization pathways into *A. baylyi* ADP1. (A) Schematic representation of the novel methodology for introducing heterologous genes for non-native substrate catabolism. (B) Comparison of strain growth on D-glucose, D-xylose, and L-arabinose as sole carbon sources. ASA547 contains the pentose catabolic gene cluster I, ASA548 contains cluster II, and ASA549 contains both clusters. Data represent average values ± standard deviations of two independent biological experiments.

Using this method, we integrated the two pentose catabolic gene clusters from *A. baumannii* into the genome of *A. baylyi* ADP1. The cluster I (∼6.6 kb), responsible for L-arabinose catabolism, was amplified by PCR and flanked with ∼1kb homologous sequences on both ends via overlap-extension PCR. The resulting linear targeting DNA fragment was designed to replace the native lactate utilization gene cluster in *A. baylyi* ADP1. Transformation was performed by adding the targeting vector directly to a culture containing mineral salts medium (MSM) supplemented with 0.8% (w/v) casein amino acids and 27 mM L-arabinose, followed by incubation. A control culture without DNA addition was also prepared. The culture with the targeting vector exhibited a significantly higher final optical density (OD) compared to the control, indicating successful enrichment of transformants. A single isolate, designated as ASA547, was obtained by further enrichment on L-arabinose as the sole carbon source and subsequent isolation by plating.

Similarly, the cluster II (∼ 9.2 kb), was integrated into the genomes of both the wild-type strain and ASA547 using D-xylose as the selective carbon source. The native butanediol utilization gene cluster was used as a neutral site for integration. The resulting strains were designated as ASA548 (harboring the cluster II) and ASA549 (harboring both the cluster I and the cluster II). Growth studies in well plates confirmed the pentose utilization capabilities of the constructed strains: ASA547 and ASA548 grew on L-arabinose and D-xylose as sole carbon sources, respectively, while ASA549 demonstrated growth on both substrates (Figure 2B). These observations align with previous reports, further validating the roles of the cluster I and the cluster II in L-arabinose and D-xylose catabolism, respectively (Alberti et al., 2023). Notably, the introduction of heterologous gene clusters did not impair host metabolism, as evidenced by the similar growth of the engineered strains and the wild-type strain on glucose.

This novel methodology represents a highly efficient strategy for constructing heterologous catabolic pathways in *A. baylyi* ADP1. By enabling transformant enrichment based on metabolic capability, the approach eliminates the need for selection markers, reducing metabolic burdens and simplifying the construction. Notably, even large gene clusters (e.g., 11 kb with flanking regions) can be efficiently integrated. This streamlined process has the potential to accelerate metabolic engineering efforts and expand the range of substrates that can be efficiently utilized by engineered strains.

### 2.3 Characterization of pentose utilization by the engineered strain

The engineered strain ASA549, harboring both pentose catabolic gene clusters, was further cultivated to characterize its growth on different sugars as sole carbon sources, as shown in Figure 3A. The strain exhibited a maximum specific growth rate of approximately 0.73 h^-1^ on D-xylose, comparable to that of the native pentose-utilizing strain *A. baumannii* (Alberti et al., 2023) and being significantly higher than ADP1’s growth rate on the native carbon source D-glucose (∼ 0.31 h^-1^). The growth rate on L-arabinose was lower (∼ 0.35 h^-1^) compared to that of D-xylose, consistent with the slower L-arabinose consumption rate. The maximum specific growth rate on D-xylose achieved here surpasses those reported for other engineered hosts, including *E. coli*, *C. glutamicum*, and *P. putida* (Table S3). Notably, growth on both D-xylose and L-arabinose showed a gradual decline in specific growth rate following the exponential phase, unlike the growth profile observed with D-glucose. Given that the pentose catabolic pathway genes in the engineered strain are induced by their respective substrates, this decline may result from the substrate-induced regulatory mechanism, where decreasing pentose concentrations reduce the induction for catabolic gene expression.

**Figure 3.**
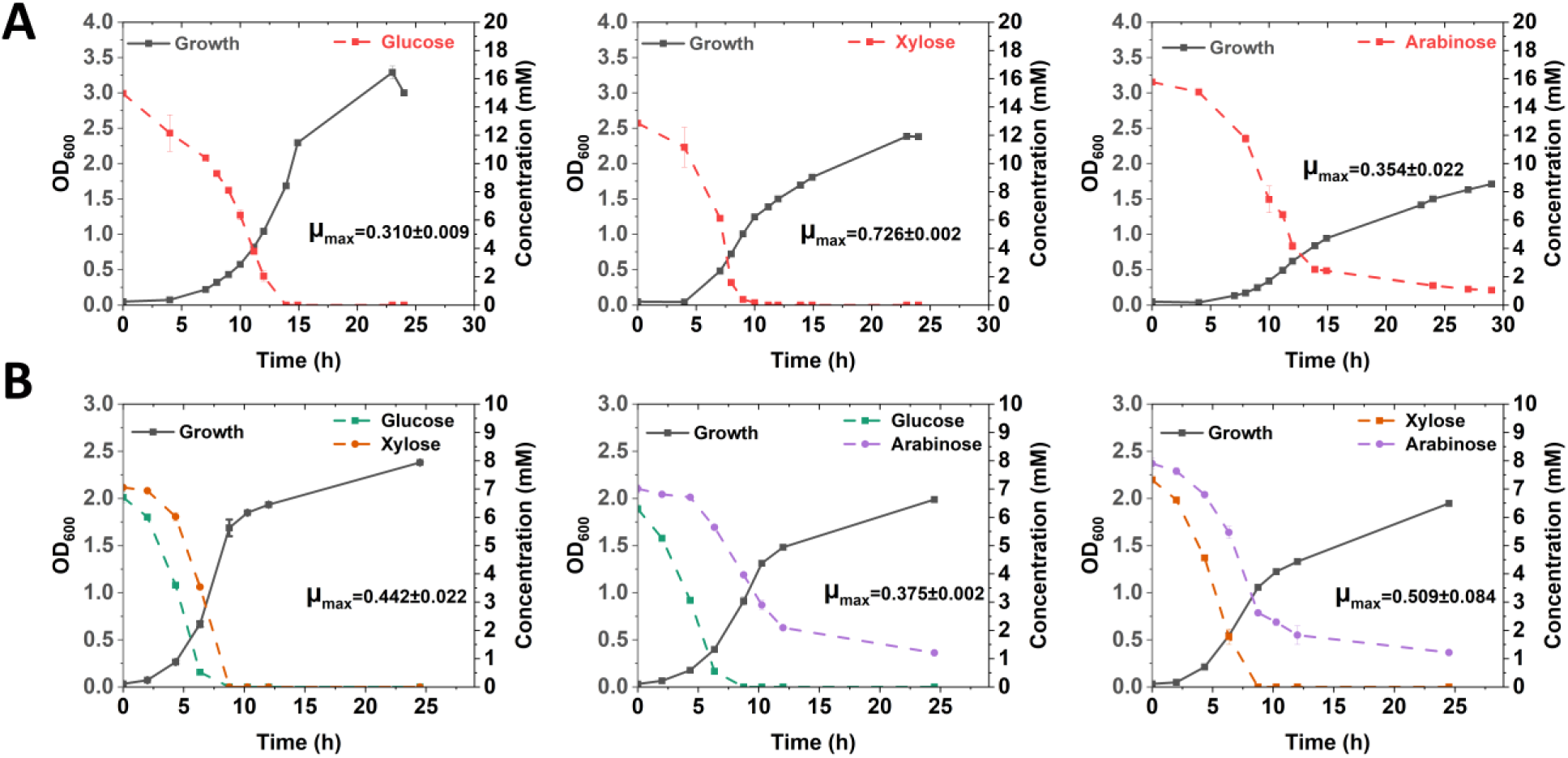
Characterization of strain ASA549 for the utilization of different sugars. (A) Growth (OD_600_) and substrate consumption when ∼15 mM D-glucose, D-xylose, or L-arabinose was used as the sole carbon source. (B) Growth and substrate consumption when sugars were used in combination (∼ 7.5 mM of each). Cells were pre-cultivated in modified lysogeny broth (LB) medium and transferred to flasks containing MSM supplemented with different substrates. The maximum specific growth rates (µ_max_, h^-1^) under each condition are presented. Data represent average values ± standard deviations of two independent biological experiments.

Interestingly, no xylonate accumulation was observed when ∼15 mM D-xylose was used as the sole carbon source, with final xylonate concentrations below 0.3 mM (Figure S2). This suggests optimal expression of downstream xylonate catabolic enzymes. The accumulation of the toxic intermediate xylonate in the Weimberg pathway has been suggested to have negative effects (Borgström et al., 2019; Meijnen et al., 2009; Rossoni et al., 2018). The absence of xylonate accumulation here highlights the balanced functionality of the engineered pathway in ASA549.

Efficient co-utilization of hemicellulose-derived sugars is critical for maximizing productivity and yield in microbial bioprocesses. Native pentose-utilizing microorganisms often experience glucose-induced catabolite repression, where pentose metabolism is repressed in the presence of D-glucose (Kim et al., 2010). We hypothesized that this repression is absent in ASA549 due to the heterologous nature of the pentose catabolic pathways. Figure 3B shows that D-xylose was consumed in the presence of D-glucose, while L-arabinose and D-xylose were simultaneously metabolized. However, a clear consumption hierarchy emerged: D-glucose > D-xylose > L- arabinose. The presence of D-glucose seemed to inhibit L-arabinose consumption. This difference in sugar consumption rates may be explained by the differences in the catalytic efficiency of the native membrane-bound Gcd (Kannisto et al., 2015). Although the Gcd in *A. baylyi* ADP1 has not been characterized, the homologous enzyme in *A. calcoaceticus*, which shares ∼ 85% identity, exhibits higher catalytic efficiency toward D-glucose than D-xylose (Dokter et al., 1987).

D-glucose, D-xylose, and L-arabinose are oxidized into their corresponding sugar acids by the membrane-bound Gcd in the periplasm before transport and catabolism. To further investigate co-utilization of sugar acids, ASA549 was cultivated on D-gluconate and D-xylonate. Interestingly, D-xylonate was consumed faster than D-gluconate (Figure S3), which is an unexpected result given that D-gluconate catabolism is native, whereas D-xylonate catabolism is non-native. Since D-xylonate is catabolized into the TCA cycle intermediate α-ketoglutarate, which can be rapidly utilized for energy production, cells may prioritize the consumption of such intermediates due to their immediate metabolic utility. However, the possibility of catabolite repression by TCA cycle intermediates cannot be ruled out.

We next investigated the metabolism of D-xylose in *A. baylyi* ASA549 by analyzing the distribution of carbon fluxes using 13C metabolic flux analysis (MFA) (Figure 4A). To our knowledge, this is the first MFA study conducted on *A. baylyi* ADP1. Here, MFA was performed with [1,2-^13^C] D-xylose. Since D-xylose catabolism directly feeds into the TCA cycle intermediate α-ketoglutarate, the 2-carbon and 3-carbon molecules are synthesized from malate and oxaloacetate via malic enzymes and phosphoenolpyruvate carboxykinase. However, the fluxes through these two pathways were not fully resolved in the current MFA experiment. Notably, 21% of the flux was directed towards acetyl-CoA from pyruvate, with only a small proportion of carbon (7.4%) entering the TCA cycle. This may be attributed to the abundant direct supply of α- ketoglutarate from D-xylose and the increased availability of reducing power generated during D-xylose oxidation. A very low carbon flow was observed through the glyoxylate shunt. To better understand the optimal fluxes during the growth on D-xylose, FBA was performed with the biomass formation reaction as the objective function (Figure 4B). The fluxes calculated by FBA were in close agreement with those determined by MFA, suggesting an optimal D-xylose metabolism in ASA549. This also highlights the advantage of the Weimberg pathway which bypasses the tightly regulated pentose phosphate pathway. In contrast, conventional pentose catabolic pathways that rely on the pentose phosphate pathway often require additional optimization or adaptation (Bator et al., 2020; Dvořák et al., 2024; Elmore et al., 2020b).

**Figure 4.**
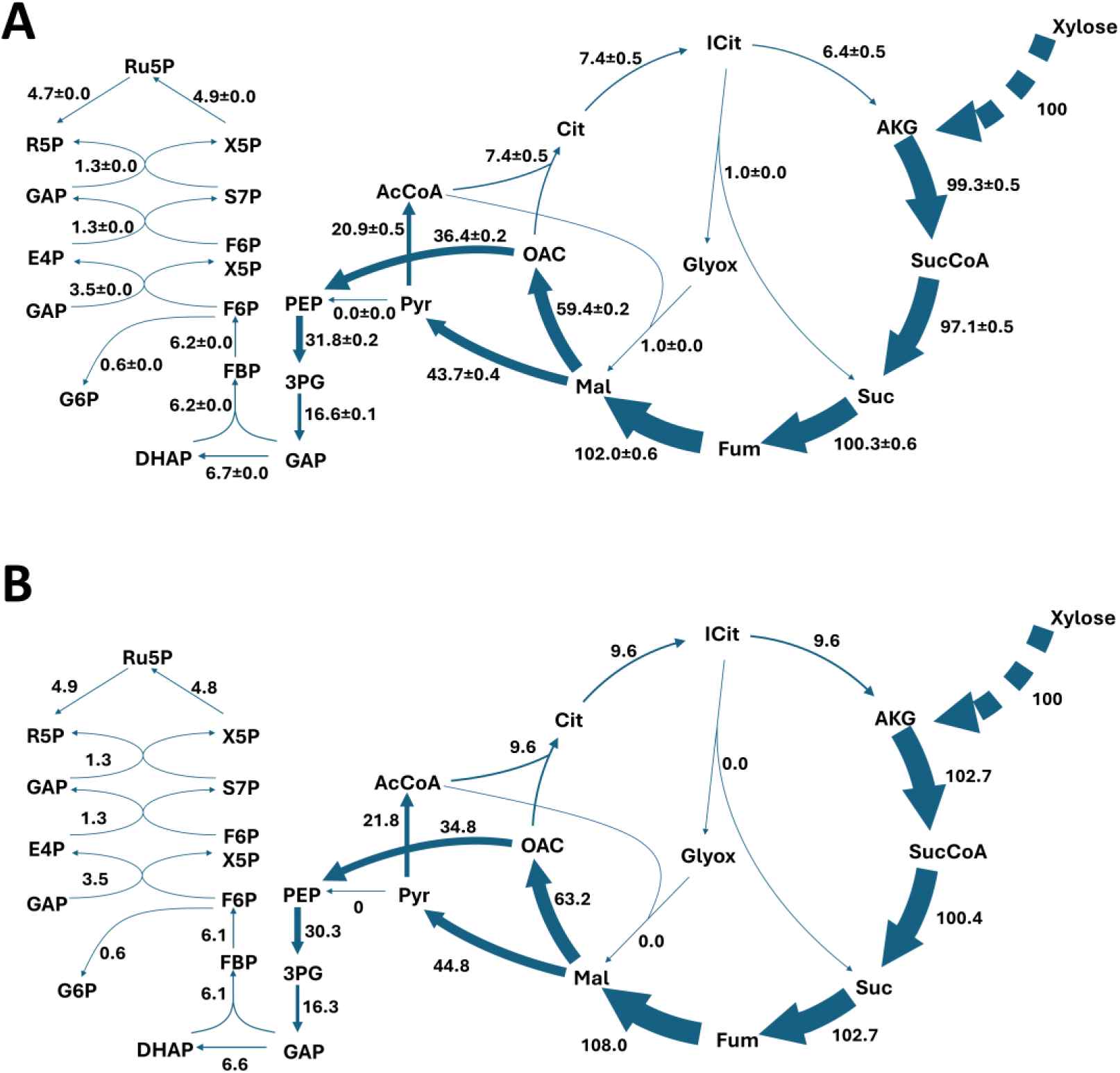
Analysis of xylose metabolism in ASA549. (A) Carbon flux distribution on D-xylose determined by ^13^C metabolic flux analysis (MFA). (B) Theoretical carbon flux distribution calculated by flux balance analysis (FBA). MFA was conducted using [1,2-^13^C] D-xylose, with flux values normalized to a specific D-xylose uptake rate of 100 mM/g CDW/h. Flux values represent average values ± standard deviations of two independent labelling experiments. The complete MFA result (including lower and upper 95% confidence interval limits) and mass isotopomer distribution data are available in Supplemental Materials S3 and S4. FBA was performed using biomass formation as the objective function. Dashed arrows indicate multiple steps. The thickness of the arrows roughly corresponds to the flux values. Abbreviations: Ru5P, ribulose 5-phosphate; R5P, ribose 5-phosphate; X5P, xylulose 5-phosphate, S7P, sedoheptulose 7- phosphate; E4P, erythrose 4-phosphate; F6B, fructose 6-phosphate; FBP, fructose 1,6- bisphosphate; GAP, glyceraldehyde3-phosphate; DHAP, dihydroxyacetone phosphate; 3PG, 3- phosphoglycerate; PEP, phosphoenolpyruvate; Pyr, pyruvate; AcCoA, acetyl-CoA; OAC, oxaloacetate; Cit, citrate; iCit, isocitrate; AKG, α-ketoglutarate; SucCoA, succinyl-CoA; Suc, succinate; Fum, fumarate; Mal, malate; Glyox, glyoxylate.

### 2.4 Demonstrating the production from D-xylose

#### 2.4.1 Engineering to redirect the carbon flow through *α*-ketoglutarate

The Weimberg pathway offers significant advantages for producing TCA cycle-derived compounds by enabling the high-yield conversion of pentoses to α-ketoglutarate while bypassing central metabolic bottlenecks. α-Ketoglutarate serves as an important precursor for the synthesis of numerous valuable compounds, including other proteinogenic amino acids (L-glutamate, L-proline, L-glutamine, and L-arginine), non-proteinogenic amino acids (GABA and L-ornithine), and various polyamide monomers (Chae et al., 2017; Lee et al., 2019). However, competition for carbon flux at the α-ketoglutarate node of the TCA cycle limits its availability for biosynthesis. Further engineering to restrict TCA cycle activity could reduce carbon loss and enhance chemical production.

To demonstrate the potential of production from α-ketoglutarate, we selected L-glutamate as the target product. To enhance L-glutamate synthesis, the native gene *gdhA* (ACIAD1110), encoding glutamate dehydrogenase, was overexpressed in ASA549 under the strong Trc promoter (Biggs et al., 2020), resulting in the strain ASA550. When cultivated in MSM supplemented with 0.2% (w/v) casein amino acids and 133 mM D-xylose, ASA550 produced ∼0.16 mM L-glutamate after 24 hours, only a ∼1.1-fold increase over ASA549. To further enhance glutamate dehydrogenase activity, the *gdhA* gene from *E. coli* was cloned into the plasmid pBAV1C-chn (Luo et al., 2019) under the control of a cyclohexanone-inducible promoter and introduced into ASA549, resulting in the strain ASA551. The plasmid has a copy number of ∼ 58 in *A. baylyi* ADP1 (Bryksin and Matsumura, 2010). Under similar conditions, ASA551 produced ∼ 0.32 mM L-glutamate after 24 hours, representing a ∼1.5-fold improvement over ASA549. However, the yield was low due to competition with cellular metabolism, particularly the demand for α-ketoglutarate in the TCA cycle.

To mitigate this competition, we hypothesized that deletion of the gene *sucAB* (ACIAD2876 and ACIAD2875), which encode the α-ketoglutarate dehydrogenase, could divert α-ketoglutarate toward L-glutamate biosynthesis (Figure 5A). The modification was also expected lead to L-glutamate accumulation, given that glutamate decarboxylase gene has not been identified in *A. baylyi* ADP1 (Durot et al., 2008) (Figure 5A). To test this, *sucAB* was deleted in ASA549, resulting in the strain ASA552. As expected, ASA552 could not grow when D-xylose was the sole carbon source (Figure S4). Interestingly, despite the *sucAB* deletion, ASA552 exhibited similar growth to ASA549 on D-glucose but exhibited a much lower final OD on succinate (Figure S4).

**Figure 5.**
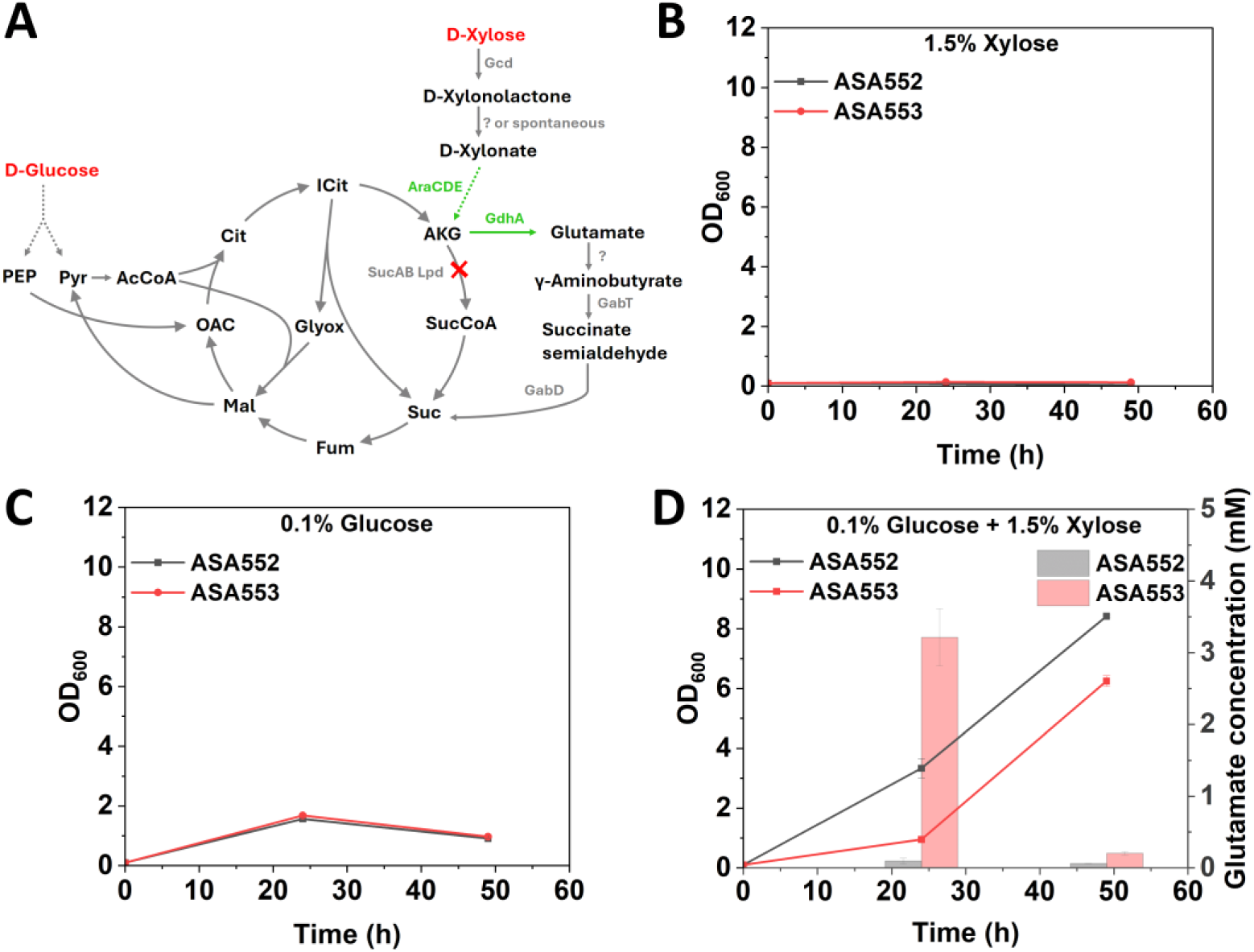
Production of L-glutamate using *sucAB* deletion strains. (A) Schematic overview of L- glutamate synthesis from D-xylose in *sucAB* deletion strains. Dashed arrows indicate pathways involving multiple steps. Heterologously expressed pathways are shown in green, and substrates used for production are highlighted in red. Flask cultivations were conducted with ASA552 and ASA553 in MSM supplemented with (B) 100 mM D-xylose, (C) 5.6 mM D-glucose, or (D) a combination of 100 mM D-xylose and 5.6 mM D-glucose. The expression of *gdhA* was induced at the beginning using 5 µM cyclohexanone. Cells were pre-cultivated in modified LB medium. Abbreviations: PEP, phosphoenolpyruvate; Pyr, pyruvate; AcCoA, acetyl-CoA; OAC, oxaloacetate; Cit, citrate; iCit, isocitrate; AKG, α-ketoglutarate; SucCoA, succinyl-CoA; Suc, succinate; Fum, fumarate; Mal, malate; Glyox, glyoxylate. Data represent average values ± standard deviations of two independent biological experiments.

Since ASA552 could not utilize D-xylose alone, a co-utilization strategy was implemented by supplementing cultures with both D-glucose and D-xylose. In this strategy, D-glucose supported cell growth while D-xylose was used for L-glutamate production. This approach is industrially relevant, as hemicellulosic hydrolysates commonly contain both pentoses and substrates such as D-glucose and lignin-derived aromatics that are utilizable by *A. baylyi* ADP1. The plasmid carrying *E. coli gdhA* was introduced into ASA552 to create ASA553. Flask cultivations of ASA552 and ASA553 were conducted. Neither strain showed growth when 100 mM D-xylose was used as the sole carbon source (Figure 5B). On 5.6 mM D-glucose, both strains could reach ODs of 1.5-1.7 (Figure 5C). Surprisingly, when 5.6 mM D-glucose and 100 mM D-xylose were combined, ASA552 and ASA553 achieved ODs of 8.4 and 6.3, respectively (Figure 5D), far exceeding their growth on D-glucose alone. In this condition, ASA553 produced ∼3.2 mM L- glutamate after 24 hours, compared to only ∼0.1 mM in ASA552 (Figure 5D), highlighting the effect of *gdhA* overexpression. However, as cell growth continued, the L-glutamate concentration decreased drastically to ∼0.2 mM after 48 h.

These findings reveal that *sucAB* deletion alone does not lead to L-glutamate accumulation. Although the *sucAB*-deficient strains could not grow on D-xylose as a sole carbon source, the presence of D-glucose facilitated D-xylose utilization for growth, as evidenced by significantly higher ODs with both carbon sources present. Although glutamate decarboxylase gene has not been identified previously in *A. baylyi* ADP1, the presence of other GABA shunt genes, *gabT* and *gabD* (Durot et al., 2008), suggests an active GABA shunt in the presence of D-glucose. This pathway likely channels L-glutamate into the TCA cycle in the *sucAB*-deficient strain (Figure 5A) and implies the existence of an unidentified glutamate decarboxylase catalyzing the conversion of L-glutamate to GABA. This hypothesis is further supported by the detection of GABA in the *A. baylyi* ADP1 metabolome (Stuani et al., 2014). Blocking the bypass pathway is not only essential for L-glutamate synthesis but also crucial for the production of a wide range of compounds derived from L-glutamate and GABA (Lee et al., 2019).

#### 2.4.2 Blocking the GABA shunt promotes L-glutamate production from *α*-ketoglutarate

We next sought to determine whether the deletion of *gabT* could prevent L-glutamate degradation and increase product yield (Figure 6A). To achieve this, the strain ASA556 was constructed by deleting *gabT* in ASA552 and introducing the plasmid harboring the *gdhA* gene from *E. coli*. Interestingly, ASA556 was unable to grow on D-glucose as the sole carbon source, likely due to a complete blockage of the TCA cycle. This finding implies the presence of an active GABA shunt in *A. baylyi* ADP1.

**Figure 6.**
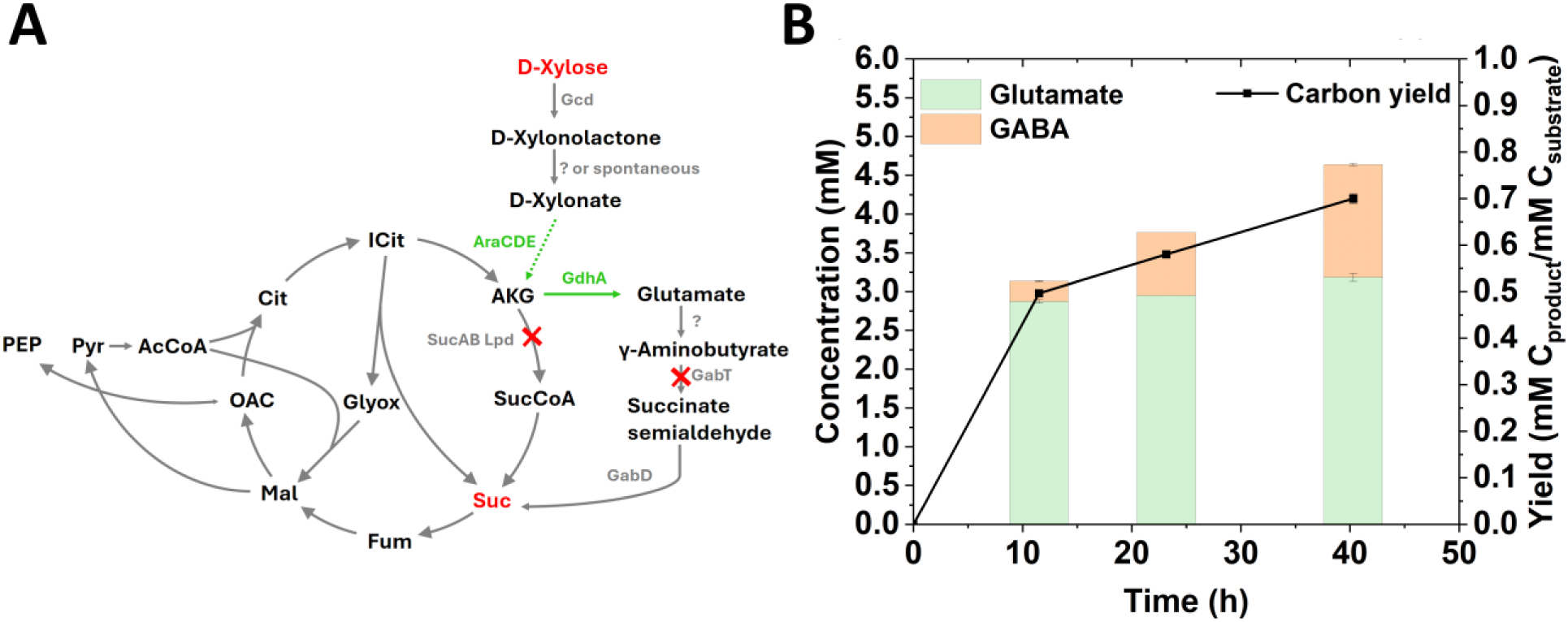
Production of L-glutamate and GABA using the *sucAB* and *gabT* deletion strain. (A) Schematic overview of L-glutamate and GABA synthesis from D-xylose in the *sucAB* and *gabT* deletion strain. Dashed arrows indicate pathways involving multiple steps. Heterologously expressed pathways are shown in green, and substrates used for production are highlighted in red. (B) L-glutamate and GABA production through whole-cell catalysis. ASA556 was initially cultivated in 40 mM succinate, with 5 µM cyclohexanone and 67 mM D-xylose added to induce *gdhA* expression and activate pentose catabolic genes. The cell pellet was then washed and resuspended in MSM supplemented with 6.7 mM D-xylose for conversion. Abbreviations: PEP, phosphoenolpyruvate; Pyr, pyruvate; AcCoA, acetyl-CoA; OAC, oxaloacetate; Cit, citrate; iCit, isocitrate; AKG, α-ketoglutarate; SucCoA, succinyl-CoA; Suc, succinate; Fum, fumarate; Mal, malate; Glyox, glyoxylate. Data represent average values ± standard deviations of two independent biological experiments.

As a proof-of-concept for high-yield D-xylose conversion, D-glucose was replaced with succinate for the co-utilization strategy. ASA556 was cultivated in MSM supplemented with 30 mM succinate and 13.3 mM D-xylose (Figure S5). After 25 hours, approximately 9.5 mM L-glutamate was produced, alongside ∼2.1 mM GABA. Despite a decrease in OD after 51 hours, the concentrations of L-glutamate and GABA increased to ∼9.9 mM and ∼2.7 mM, respectively, representing a total carbon yield of ∼34 % C_product_/C_xylose+succinate_. The production of GABA confirmed the presence of an unidentified enzyme for L-glutamate decarboxylation. The glutamate decarboxylase system has evolved in several bacterial genera as a defense mechanism in acidic environments, utilizing protons and glutamate as substrates for GABA production (Feehily and Karatzas, 2013). In organisms like E*. coli*, this system functions optimally under acidic conditions and loses activity near neutral pH (Guan et al., 2025). The unidentified mechanism responsible for GABA production in *A. baylyi* ADP1 may indicate a distinct regulation.

To further assess L-glutamate and GABA production solely from D-xylose, whole-cell catalysis using resting cells was performed (Figure 6B). Cells were first grown on succinate and then washed and resuspended in MSM containing 6.7 mM D-xylose. With a conversion OD of ∼1.5, L-glutamate and GABA concentrations reached ∼2.9 mM and ∼0.26 mM, respectively, after 11 hours.

After 40 hours, these concentrations increased to ∼3.2 mM and ∼1.5 mM, yielding a total carbon conversion of ∼70 % C_product_/C_xylose_. Approximately 1.8 mM xylonate remained in the medium, representing the residual carbon. The incomplete conversion could be attributed to insufficient biomass or imbalance of energy/cofactors. A similar synthetic pathway was previously implemented in *E. coli*, where the Weimberg pathway was introduced for GABA production, and native D-xylose metabolism was blocked (Zhao et al., 2017). In that study, cell growth was supported with yeast extract and tryptone, while D-xylose was utilized for L-glutamate and GABA production. The total carbon yield of L-glutamate and GABA was estimated at ∼0.4 mM C _product_/mM C _xylose_, which was lower than the yield achieved in our study.

Since the deletion of both *sucAB* and *gabT* makes the strain unable to grow on D-glucose, restoring growth on this substrate is of industrial significance, as D-glucose is an important component of cellulosic and hemicellulosic hydrolysates. To explore metabolic flux alterations compared to the wild-type strain, we performed FBA on the double deletion strain with biomass formation as the objective. The FBA predictions indicated that the deletion of *sucAB* and *gabT* was not lethal. Significant metabolic changes included upregulation of the glyoxylate shunt, malic enzyme reactions, and the pyruvate dehydrogenase reaction (Figure S6), which collectively generate substantial amounts of NAD(P)H to compensate for the disrupted TCA cycle. Future efforts will involve the use of adaptive laboratory evolution (ALE) to enable growth of the *sucAB* and *gabT* deletion strain on D-glucose, thereby resolving this bottleneck. ALE provides the advantage of generating mutants with multiple beneficial mutations, including those that are non-intuitive. Notably, ALE has successfully restored growth in a TCA cycle-deficient *E. coli* strain cultivated on glucose minimal medium (Santala et al., 2011; Zhou et al., 2024). Given the genome plasticity of *A. baylyi* ADP1, its natural competence, and high recombination activity, this organism is particularly well-suited for ALE and identifying advantageous mutations through reverse engineering (Luo et al., 2022).

## 3. Conclusions

In this study, we successfully engineered *A. baylyi* ADP1 to utilize D-xylose and L-arabinose by introducing the heterologous Weimberg pathway from *A. baumannii*. Our marker-free approach enabled efficient integration of the two pentose catabolic gene clusters into the genome. Growth studies and metabolic flux analysis validated the optimized metabolism, highlighting the advantages of the Weimberg pathway in bypassing key regulatory bottlenecks.

To explore the potential for high-value bioproduct synthesis, L-glutamate was selected as a target compound. While strategies such as *gdhA* overexpression and *sucAB* deletion demonstrated limited success due to persistent L-glutamate degradation, likely mediated by an unidentified glutamate decarboxylase, the subsequent deletion of *gabT* produced a TCA-deficient strain capable of producing L-glutamate and GABA at high yields, though with impaired growth on D-glucose.

This study not only establishes an efficient platform for pentose utilization but also demonstrates the feasibility of producing TCA-derived compounds using engineered *A. baylyi* strains. Future work could focus on identifying metabolic bottlenecks for growth recovery and optimizing processes for scalable bioproduction.

## 4. Experimental procedures

### 4.1 Strains and media

*E. coli* XL1-Blue (Stratagene, USA) was used for molecular cloning work. *A. baylyi* ADP1 (DSM 24193, DSMZ, Germany) was used as the host for all the experiments. All the strains used in the study are listed in Table 1.

**Table 1.** Strains and plasmids used in the study.

| Strain name | Organism | Genotype or description | Source or reference |
| --- | --- | --- | --- |
| XL1-Blue | *E. coli* | *recA1 endA1 gyrA96 thi-1 hsdR17 supE44 relA1 lac* [F' *proAB lacIqZΔM15 Tn10 Tet<sup>r</sup>*] | Stratagene, USA |
| WT ADP1 | *A. baylyi* ADP1 | Wild-type | DSM 24193, DSMZ, Germany |
| ASA546 | *A. baylyi* ADP1 | WT ADP1 carrying pJL19 | This study |
| ASA547 | *A. baylyi* ADP1 | *ΔlldP-dld::ara* gene cluster I | This study |
| ASA548 | *A. baylyi* ADP1 | *ΔacoA-budC::ara* gene cluster II | This study |
| ASA549 | *A. baylyi* ADP1 | *ΔlldP-dld::ara* gene cluster I, *ΔacoA-budC::ara* gene cluster II | This study |
| ASA550 | *A. baylyi* ADP1 | *ΔgdhA* (ACIAD1110)::*tdk-kan<sup>r</sup>*-P<sub>trc</sub>-*gdhA* (ACIAD1110) | This study |
| ASA551 | *A. baylyi* ADP1 | ASA549 carrying pJL20 | This study |
| ASA552 | *A. baylyi* ADP1 | *ΔlldP-dld::ara* gene cluster I, *ΔacoA-budC::ara* gene cluster II, *ΔsucAB::tdk-kan<sup>r</sup>* | This study |
| ASA553 | *A. baylyi* ADP1 | ASA552 carrying pJL20 | This study |
| ASA554 | | *lldP-dld::ara* gene cluster I, Δ*acoA-budC::ara* gene cluster II, Δ*sucAB* | This study |
| ASA555 | *A. baylyi* ADP1 | Δ*lldP-dld::ara* gene cluster I, Δ*acoA-budC::ara* gene cluster II, Δ*sucAB*, Δ*gabT::tdk-kan<sup>r</sup>* | This study |
| ASA556 | *A. baylyi* ADP1 | ASA555 carrying pJL20 | This study |

Modified lysogeny broth (LB) medium (10 g/L tryptone, 5 g/L yeast extract, and 1 g/L NaCl) was used to cultivate *E. coli* and *A. baylyi* ADP1 during cloning and genetic manipulation. When required, 22 mM D-glucose was added to the medium. For *A*. *baylyi* ADP1 strains with a deletion of the *sucAB* and *gabT* genes, growth was supported using MSM supplemented with 25 mM succinate, as these strains exhibited poor growth in LB medium. MSM was used for growth studies and production experiments, with substrates including, D-glucose/sodium D-gluconate, D- xylose/sodium D-xylonate, L-arabinose, succinate, and casein amino acids added as specified in the result and discussion section. The composition of MSM is 3.88 g/L K_2_HPO_4_, 1.63 g/L NaH_2_PO_4_, 2.00 g/L (NH4)_2_SO_4_, 0.1 g/L MgCl_2_·6H_2_O, 10 mg/L ethylenediaminetetraacetic acid (EDTA), 2 mg/L ZnSO_4_·7H_2_O, 1 mg/L CaCl_2_·2H_2_O, 5 mg/L FeSO_4_·7H_2_O, 0.2 mg/L Na_2_MoO_4_·2H_2_O, 0.2 mg/L CuSO_4_·5H_2_O, 0.4 mg/L CoCl_2_·6H_2_O, and 1 mg/L MnCl_2_·2H_2_O. Antibiotics were added when required (25 μg/ml chloramphenicol and 50 μg/ml kanamycin).

### 4.2 Culture conditions

Cells were pre-cultivated overnight in modified LB medium and subsequently inoculated into MSM supplemented with appropriate carbon sources for growth studies to characterize pentose utilization. For strains with *sucAB* deletion, precultivation was conducted in MSM containing 25 mM succinate. In well-plate cultivation, cells were grown in a 200 µL volume using 96-well plates (flat bottom, μClear™, white, Greiner) at 30°C in a Spark multimode microplate reader (Tecan, Switzerland). Double orbital shaking (6 mm amplitude, 54 rpm) was performed for 5 minutes twice per hour, and OD at 600 nm was measured hourly. Flask cultivation was carried out in 250 mL flasks containing a 50 mL culture volume, incubated at 30°C and 300 rpm.

For L-glutamate and GABA production, cells were pre-cultivated in MSM supplemented with 25 mM succinate, followed by cultivation in 100 mL flasks containing a 10 mL culture volume at 30°C and 300 rpm, with MSM supplemented with appropriate carbon sources. When induction was required, 5 µM cyclohexanone was added from the beginning of cultures. Whole-cell catalysis involved initial cultivation in MSM containing 40 mM succinate, 5 µM cyclohexanone, and 67 mM D-xylose. After overnight cultivation, cells were washed and resuspended in MSM supplemented with 6.7 mM D-xylose, and the conversion was carried out in 14 mL cultivation tubes at 30°C and 300 rpm.

### 4.3 Genetic engineering

Molecular cloning was performed following standard protocols. Transformation and the homologous recombination-based genome editing of *A. baylyi* ADP1 were carried out as described previously (Santala et al., 2011). The plasmids used in this study are listed in Table 2 and all the primers used in this study along with detailed descriptions can be found in Supplemental materials S2.

**Table 2.** Plasmids used in the study.

| Plasmid name | Description | Source or reference |
| --- | --- | --- |
| pBAV1C-chn | broad-host range vector, pBAV ori, chnR, P<sub>chnB</sub>, Cm<sup>r</sup> | (Luo et al., 2019) |
| pJL19 | pBAV1C-chn carrying *araD1* (HTZ92_1391) from *A. baumannii* | This study |
| pJL20 | pBAV1C-chn carrying *gdhA* from *E. coli* BW25113 | This study |

Strains ASA547, ASA548, and ASA549 were constructed by integrating pentose catabolic gene clusters using a novel approach based on natural transformation and growth-based selection. The pentose catabolic gene clusters were amplified from the genome of *A. baumannii* ATCC 19606 using Phusion high-fidelity DNA polymerase (Thermo Fisher Scientific, Finland). The pentose catabolic gene cluster I and II were flanked with homologous sequences targeting the lactate catabolic gene cluster (*lldP* (ACIAD0106)-*dld* (ACIAD0109)) and butanediol catabolic gene cluster (*acoA* (ACIAD1017)-*budC* (ACIAD1022)), respectively, using overlap-extension PCR. The assembled PCR products were further purified with GeneJET Gel Extraction kit (Thermo Fisher Scientific). For transformation and transformant enrichment, 10 µl of the purified DNA (∼500 ng) was added to 190 µl MSM supplemented with 0.8% (w/v) casein amino acids and 27 mM L-arabinose/D-xylose in well-plates. The medium was further inoculated with the cells to be transformed, and a culture without DNA addition was included as a control. The plate was incubated on a shaker for up to 24 hours. A significantly higher OD in cultures with DNA addition compared to the control indicated successful enrichment of pentose-utilizing transformants. Further enrichment and isolation were performed by transferring cells to fresh medium containing pentose as the sole carbon source, followed by plating for colony isolation.

The plasmid pJL19 was constructed by assembling the araD1 gene, amplified from the genome of *A. baumannii* ATCC 19606, with the linearized pBAV1C-chn plasmid (Luo et al., 2019) using Gibson Assembly. Plasmid pJL20 was constructed by restriction and ligation using SpeI and PvuI between the pBAV1C-chn plasmid and the *gdhA* gene amplified from *E. coli* BW25113. Overexpression of the native *gdhA* gene (ACIAD1110) was achieved by placing the strong Trc promoter and a strong ribosome binding site (RBS) upstream of the gene. The linear integration cassette for *gdhA* overexpression was constructed by assembling the left homologous flanking sequence, the tdk/kan cassette (Metzgar, 2004), and the right homologous flanking sequence using overlap-extension PCR; the Trc promoter and RBS sequence were included in the primer used to amplify the right homologous flanking sequence. Linear cassettes for *sucAB* and *gabT* knock-out were constructed by assembling the left homologous flanking sequence, the tdk/kan cassette (Metzgar, 2004), and the right homologous flanking sequence using overlap-extension PCR. Removal of the antibiotic marker was achieved by transformation with a rescue cassette, followed by counter-selection as previously described (Metzgar, 2004).

### 4.4 Analytical methods

Substrate and product concentrations were determined with high-performance liquid chromatography (HPLC). D-glucose, D-xylose, and L-arabinose were analyzed with an LC-20 AC prominence liquid chromatograph (Shimadzu, USA) equipped with RID-10A refractive Index detector. Separation was performed on a Phenomenex Rezex RHM-monosaccharide H⁺ (8%) column with Milli-Q water as the mobile phase at a flow rate of 0.5 mL/min and a temperature of 40°C. For D-gluconate and D-xylonate, 5 mM sulfuric acid was used as the mobile phase instead of water.

When sulfuric acid was used as the mobile phase, peaks for sugars and sugar acids overlapped. In the presence of both sugars and sugar acids, sugar acid concentrations were estimated as follows: sugar concentrations were first determined using Milli-Q water as the mobile phase. The peak areas for sugars when using 5 mM sulfuric acid as the mobile phase were estimated from a standard curve and the previously determined sugar concentrations. The sugar acid peak areas were then calculated by subtracting the estimated sugar areas from the total peak areas obtained when sulfuric acid was used as the mobile phase.

L-glutamate and GABA were derivatized prior to HPLC analysis following a previously described protocol (Zhang et al., 2020). Briefly, 50 μL of a derivatization reagent (2,4- dinitrofluorobenzene/acetonitrile, 1:99) and 100 μL of 0.5 M NaHCO_3_ were added to 100 μL of the supernatant. The mixture was incubated at 60°C for 60 minutes. The reaction was stopped by adding 750 μL of 0.01 M KH_2_PO_4_, followed by shaking and resting for 10 minutes. Samples were analyzed using an LC-40D HPLC system (Shimadzu, Japan) with a Luna 5 µm C18 column (150 × 4.6 mm, Phenomenex, USA) and a photodiode array detector (SPD-M40). The analysis was performed at 40°C using a mobile phase of 0.1% (v/v) formic acid and acetonitrile (65:35) at a flow rate of 0.5 mL/min.

### 4.5 ^13^C labelling experiments and metabolic flux analysis

The ^13^C labelling experiment was designed using the software influx_s (Sokol et al., 2012), with [1,2-^13^C] D-xylose selected as the labeling substrate. The experiment was conducted in duplicate using strain ASA549. Cells were pre-cultivated in modified LB medium for approximately 6 hours, followed by centrifugation (3000 rpm, 5 min) and washing with MSM. The pellet was then resuspended in 10 mL MSM containing 0.2% (w/v) [1,2-^13^C] D-xylose in 100 mL flasks. To minimize unlabeled biomass, the initial OD was adjusted to ∼0.008. Cultivation was carried out at 30°C and 300 rpm, with periodic sampling for OD measurement, as well as substrate and extracellular metabolite concentration analysis. Once the OD reached 0.8–0.9, samples were collected for analyzing the labeling pattern of proteinogenic amino acids.

For proteinogenic amino acid labeling analysis, the protocol described in reference was followed (Long and Antoniewicz, 2019). Briefly, 500 μL of the sample was centrifuged to collect the pellet, which was then hydrolyzed with 500 μL of 6 N HCl at 110°C for 12–18 hours to release amino acids. After hydrolysis, the sample was centrifuged to remove debris, and 400 μL of the supernatant was transferred to a new tube and evaporated at 65°C under air or nitrogen flow. The dried sample was then derivatized by adding 35 μL of pyridine and 50 μL of MTBSTFA + 1% (wt/wt) TBDMSCl (Sigma-Aldrich, cat. no. 375934), followed by incubation at 60°C for 30 minutes. Finally, 75 μL of the derivatized sample was transferred to a vial for analysis by gas chromatography-mass spectrometry (GC-MS). GC-MS was performed using a GCMS-QP2020 NX gas chromatograph-mass spectrometer (Shimadzu) with a Nexis GC-2030 system (Shimadzu) and a Zebron ZB-5MSi GC column (30 m × 0.25 mm × 0.25 µm, Phenomenex). The injection volume was set to 1 μL, with helium (1 mL/min) as the carrier gas. The temperature program was as follows: 80°C (2 min hold), ramped to 280°C at 7°C/min, and held for 20 minutes.

The isotopomer distribution data was corrected for natural isotopic abundance using IsoCor (Millard et al., 2019). The corrected isotopomer distribution data are provided in Supplemental Materials S3. Flux analysis was performed using both mfapy (Matsuda et al., 2021) and influx_s (Sokol et al., 2012), with the central metabolic model constrained by mass isotopomer distribution data of proteinogenic amino acids, as well as specific growth and substrate uptake rates. The central metabolic network of *A. baylyi* ADP1 was prepared based on its genome-scale metabolic model (Durot et al., 2008). Confidence intervals were calculated using influx_s (Sokol et al., 2012).

### 4.6 Flux balance analysis

The previously described genome-wide *A. baylyi* metabolic network (Durot et al., 2008) was used for the implementation of FBA. FBA was performed using COBRApy (Ebrahim et al., 2013). To simulate theoretical yields of different TCA cycle intermediates via different pentose utilization pathway (Table S1), different pathways are incorporated into the metabolic model (as detailed in Table S2). To predict the growth of the *sucAB* and *gabT* deletion strain on D-glucose and simulate the optimal flux distribution, the flux of the α-ketoglutarate dehydrogenase reaction was constrained to zero. For the simulations, the carbon source intake rate was set to 10 mM/h/g CDW with other nutrient unlimited.

## Acknowledgements

Funding: SS would like to thank the Novo Nordisk Foundation (grant NNF21OC0067758) and the Research Council of Finland (grant no. 347204 and 353587). VS would like to thank the Novo Nordisk Foundation (grant NNF21OC0079579). This research was supported from the European Union – NextGenerationEU instrument and is funded by the Research Council of Finland under grant number 353658.

## Author contribution

Author contribution: JL, SS, and VS designed the study. JL and EE carried out the research work. JL, SS, and VS analyzed the data. SS and VS supervised the study. All authors participated in writing the manuscript. All authors read and approved the final manuscript.

## References

Alberti, L., König, P., Zeidler, S., Poehlein, A., Daniel, R., Averhoff, B., Müller, V., 2023. Identification and characterization of a novel pathway for aldopentose degradation in Acinetobacter baumannii. Environ Microbiol 25, 2416–2430. 10.1111/1462-2920.16471

Arvay, E., Biggs, B.W., Guerrero, L., Jiang, V., Tyo, K., 2021. Engineering Acinetobacter baylyi ADP1 for mevalonate production from lignin-derived aromatic compounds. Metab Eng Commun 13, e00173. 10.1016/j.mec.2021.e00173

Barbe, V., 2004. Unique features revealed by the genome sequence of Acinetobacter sp. ADP1, a versatile and naturally transformation competent bacterium. Nucleic Acids Res 32, 5766–5779. 10.1093/nar/gkh910

Bator, I., Wittgens, A., Rosenau, F., Tiso, T., Blank, L.M., 2020. Comparison of Three Xylose Pathways in Pseudomonas putida KT2440 for the Synthesis of Valuable Products. Front Bioeng Biotechnol 7. 10.3389/fbioe.2019.00480

Beckham, G.T., Johnson, C.W., Karp, E.M., Salvachúa, D., Vardon, D.R., 2016. Opportunities and challenges in biological lignin valorization. Curr Opin Biotechnol 42, 40–53. 10.1016/j.copbio.2016.02.030

Bedore, S.R., Neidle, E.L., Pardo, I., Luo, J., Baugh, A.C., Duscent-Maitland, C. V., Tumen-Velasquez, M.P., Santala, V., Santala, S., 2023. Natural transformation as a tool in Acinetobacter baylyi: Streamlined engineering and mutational analysis. pp. 207–234. 10.1016/bs.mim.2023.01.002

Biggs, B.W., Bedore, S.R., Arvay, E., Huang, S., Subramanian, H., McIntyre, E.A., Duscent-Maitland, C. V., Neidle, E.L., Tyo, K.E.J., 2020. Development of a genetic toolset for the highly engineerable and metabolically versatile Acinetobacter baylyi ADP1. Nucleic Acids Res 48, 5169–5182. 10.1093/nar/gkaa167

Borgström, C., Wasserstrom, L., Almqvist, H., Broberg, K., Klein, B., Noack, S., Lidén, G., Gorwa-Grauslund, M.F., 2019. Identification of modifications procuring growth on xylose in recombinant Saccharomyces cerevisiae strains carrying the Weimberg pathway. Metab Eng 55, 1–11. 10.1016/j.ymben.2019.05.010

Bryksin, A. V., Matsumura, I., 2010. Rational Design of a Plasmid Origin That Replicates Efficiently in Both Gram-Positive and Gram-Negative Bacteria. PLoS One 5, e13244. 10.1371/journal.pone.0013244

Chae, T.U., Ko, Y.-S., Hwang, K.-S., Lee, S.Y., 2017. Metabolic engineering of Escherichia coli for the production of four-, five- and six-carbon lactams. Metab Eng 41, 82–91. 10.1016/j.ymben.2017.04.001

Choi, J.W., Jeon, E.J., Jeong, K.J., 2019. Recent advances in engineering Corynebacterium glutamicum for utilization of hemicellulosic biomass. Curr Opin Biotechnol 57, 17–24. 10.1016/j.copbio.2018.11.004

de Berardinis, V., Durot, M., Weissenbach, J., Salanoubat, M., 2009. Acinetobacter baylyi ADP1 as a model for metabolic system biology. Curr Opin Microbiol 12, 568–576. 10.1016/j.mib.2009.07.005

Dokter, P., Pronk, J.T., Schie, B.J., Dijken, J.P., Duine, J.A., 1987. The in vivo and in vitro substrate specificity of quinoprotein glucose dehydrogenase of *Acinetobacter calcoaceticus* LMD79.41. FEMS Microbiol Lett 43, 195–200. 10.1111/j.1574-6968.1987.tb02122.x

Durot, M., Le Fèvre, F., de Berardinis, V., Kreimeyer, A., Vallenet, D., Combe, C., Smidtas, S., Salanoubat, M., Weissenbach, J., Schachter, V., 2008. Iterative reconstruction of a global metabolic model of Acinetobacter baylyi ADP1 using high-throughput growth phenotype and gene essentiality data. BMC Syst Biol 2, 85. 10.1186/1752-0509-2-85

Dvořák, P., Burýšková, B., Popelářová, B., Ebert, B.E., Botka, T., Bujdoš, D., Sánchez-Pascuala, A., Schöttler, H., Hayen, H., de Lorenzo, V., Blank, L.M., Benešík, M., 2024. Synthetically-primed adaptation of Pseudomonas putida to a non-native substrate D-xylose. Nat Commun 15. 10.1038/s41467-024-46812-9

Ebrahim, A., Lerman, J.A., Palsson, B.O., Hyduke, D.R., 2013. COBRApy: COnstraints-Based Reconstruction and Analysis for Python. BMC Syst Biol 7, 74. 10.1186/1752-0509-7-74

Elmore, J.R., Dexter, G.N., Salvachúa, D., O’Brien, M., Klingeman, D.M., Gorday, K., Michener, J.K., Peterson, D.J., Beckham, G.T., Guss, A.M., 2020a. Engineered Pseudomonas putida simultaneously catabolizes five major components of corn stover lignocellulose: Glucose, xylose, arabinose, p- coumaric acid, and acetic acid. Metab Eng 62, 62–71. 10.1016/j.ymben.2020.08.001

Elmore, J.R., Dexter, G.N., Salvachúa, D., O’Brien, M., Klingeman, D.M., Gorday, K., Michener, J.K., Peterson, D.J., Beckham, G.T., Guss, A.M., 2020b. Engineered Pseudomonas putida simultaneously catabolizes five major components of corn stover lignocellulose: Glucose, xylose, arabinose, p- coumaric acid, and acetic acid. Metab Eng 62, 62–71. 10.1016/j.ymben.2020.08.001

Feehily, C., Karatzas, K.A.G., 2013. Role of glutamate metabolism in bacterial responses towards acid and other stresses. J Appl Microbiol 114, 11–24. 10.1111/j.1365-2672.2012.05434.x

Geng, B., Jia, X., Peng, X., Han, Y., 2022. Biosynthesis of value-added bioproducts from hemicellulose of biomass through microbial metabolic engineering. Metab Eng Commun 15, e00211. 10.1016/j.mec.2022.e00211

Guan, F., Fu, B., Wang, P., Yan, C., Wu, M., Xu, X., Wang, H., Yu, P., 2025. Directed evolution of glutamate decarboxylase B for enhancing its enzyme activity towards nearly neutral pHs based on error-prone PCR. Int J Biol Macromol 292, 139283. 10.1016/j.ijbiomac.2024.139283

Guo, H., Zhao, Y., Chang, J.-S., Lee, D.-J., 2022. Inhibitor formation and detoxification during lignocellulose biorefinery: A review. Bioresour Technol 361, 127666. 10.1016/j.biortech.2022.127666

Kannisto, M.S., Mangayil, R.K., Shrivastava-Bhattacharya, A., Pletschke, B.I., Karp, M.T., Santala, V.P., 2015. Metabolic engineering of Acinetobacter baylyi ADP1 for removal of Clostridium butyricum growth inhibitors produced from lignocellulosic hydrolysates. Biotechnol Biofuels 8, 198. 10.1186/s13068-015-0389-6

Kaplan, N.A., Islam, K.N., Kanis, F.C., Verderber, J.R., Wang, X., Jones, J.A., Koffas, M.A.G., 2024. Simultaneous glucose and xylose utilization by an Escherichia coli catabolite repression mutant. Appl Environ Microbiol 90. 10.1128/aem.02169-23

Kim, D., Woo, H.M., 2018. Deciphering bacterial xylose metabolism and metabolic engineering of industrial microorganisms for use as efficient microbial cell factories. Appl Microbiol Biotechnol. 10.1007/s00253-018-9353-2

Kim, J.-H., Block, D.E., Mills, D.A., 2010. Simultaneous consumption of pentose and hexose sugars: an optimal microbial phenotype for efficient fermentation of lignocellulosic biomass. Appl Microbiol Biotechnol 88, 1077–1085. 10.1007/s00253-010-2839-1

Kuschmierz, L., Shen, L., Bräsen, C., Snoep, J., Siebers, B., 2022. Workflows for optimization of enzyme cascades and whole cell catalysis based on enzyme kinetic characterization and pathway modelling. Curr Opin Biotechnol. 10.1016/j.copbio.2021.10.020

Lee, S.Y., Kim, H.U., Chae, T.U., Cho, J.S., Kim, J.W., Shin, J.H., Kim, D.I., Ko, Y.-S., Jang, W.D., Jang, Y.-S., 2019. A comprehensive metabolic map for production of bio-based chemicals. Nat Catal 2, 18–33. 10.1038/s41929-018-0212-4

Li, X., Chen, Y., Nielsen, J., 2019. Harnessing xylose pathways for biofuels production. Curr Opin Biotechnol. 10.1016/j.copbio.2019.01.006

Linh, T.N., Fujita, H., Sakoda, A., 2017. Release kinetics of esterified p-coumaric acid and ferulic acid from rice straw in mild alkaline solution. Bioresour Technol 232, 192–203. 10.1016/j.biortech.2017.02.009

Liu, C., Choi, B., Efimova, E., Nygård, Y., Santala, S., 2024a. Enhanced upgrading of lignocellulosic substrates by coculture of Saccharomyces cerevisiae and Acinetobacter baylyi ADP1. Biotechnology for Biofuels and Bioproducts 17, 61. 10.1186/s13068-024-02510-8

Liu, C., Efimova, E., Santala, V., Santala, S., 2024b. Analysis of detoxification kinetics and end products of furan aldehydes in Acinetobacter baylyi ADP1. Sci Rep 14, 29678. 10.1038/s41598-024-81124-4

Long, B., Zhang, F., Dai, S.Y., Foston, M., Tang, Y.J., Yuan, J.S., 2024. Engineering strategies to optimize lignocellulosic biorefineries. Nature Reviews Bioengineering. 10.1038/s44222-024-00247-5

Long, C.P., Antoniewicz, M.R., 2019. High-resolution 13C metabolic flux analysis. Nat Protoc 14, 2856– 2877. 10.1038/s41596-019-0204-0

Luo, J., Lehtinen, T., Efimova, E., Santala, V., Santala, S., 2019. Synthetic metabolic pathway for the production of 1-alkenes from lignin-derived molecules. Microb Cell Fact 18, 48. 10.1186/s12934-019-1097-x

Luo, J., McIntyre, E.A., Bedore, S.R., Santala, V., Neidle, E.L., Santala, S., 2022. Characterization of Highly Ferulate-Tolerant Acinetobacter baylyi ADP1 Isolates by a Rapid Reverse Engineering Method. Appl Environ Microbiol 88. 10.1128/AEM.01780-21

Matsuda, F., Maeda, K., Taniguchi, T., Kondo, Y., Yatabe, F., Okahashi, N., Shimizu, H., 2021. mfapy: An open-source Python package for 13C-based metabolic flux analysis. Metab Eng Commun 13, e00177. 10.1016/j.mec.2021.e00177

Meijnen, J.-P., de Winde, J.H., Ruijssenaars, H.J., 2009. Establishment of Oxidative <SCP>d</SCP>-Xylose Metabolism in *Pseudomonas putida* S12. Appl Environ Microbiol 75, 2784–2791. 10.1128/AEM.02713-08

Meriläinen, E., Efimova, E., Santala, V., Santala, S., 2024. Carbon-wise utilization of lignin-related compounds by synergistically employing anaerobic and aerobic bacteria. Biotechnology for Biofuels and Bioproducts 17, 78. 10.1186/s13068-024-02526-0

Metzgar, D., 2004. Acinetobacter sp. ADP1: an ideal model organism for genetic analysis and genome engineering. Nucleic Acids Res 32, 5780–5790. 10.1093/nar/gkh881

Millard, P., Delépine, B., Guionnet, M., Heuillet, M., Bellvert, F., Létisse, F., 2019. IsoCor: isotope correction for high-resolution MS labeling experiments. Bioinformatics 35, 4484–4487. 10.1093/bioinformatics/btz209

Mills, T.Y., Sandoval, N.R., Gill, R.T., 2009. Cellulosic hydrolysate toxicity and tolerance mechanisms in Escherichia coli. Biotechnol Biofuels 2, 26. 10.1186/1754-6834-2-26

Narisetty, V., Cox, R., Bommareddy, R., Agrawal, D., Ahmad, E., Pant, K.K., Chandel, A.K., Bhatia, S.K., Kumar, D., Binod, P., Gupta, V.K., Kumar, V., 2022. Valorisation of xylose to renewable fuels and chemicals, an essential step in augmenting the commercial viability of lignocellulosic biorefineries. Sustain Energy Fuels 6, 29–65. 10.1039/D1SE00927C

Qiu, Y., Wu, M., Bao, H., Liu, W., Shen, Y., 2023. Engineering of Saccharomyces cerevisiae for co-fermentation of glucose and xylose: Current state and perspectives. Engineering Microbiology. 10.1016/j.engmic.2023.100084

Rossoni, L., Carr, R., Baxter, S., Cortis, R., Thorpe, T., Eastham, G., Stephens, G., 2018. Engineering Escherichia coli to grow constitutively on D-xylose using the carbon-efficient Weimberg pathway. Microbiology (N Y) 164, 287–298. 10.1099/mic.0.000611

Salmela, M., Lehtinen, T., Efimova, E., Santala, S., Santala, V., 2019. Alkane and wax ester production from lignin-related aromatic compounds. Biotechnol Bioeng 116, 1934–1945. 10.1002/bit.27005

Salvachúa, D., Karp, E.M., Nimlos, C.T., Vardon, D.R., Beckham, G.T., 2015. Towards lignin consolidated bioprocessing: simultaneous lignin depolymerization and product generation by bacteria. Green Chemistry 17, 4951–4967. 10.1039/C5GC01165E

Santala, S., Efimova, E., Kivinen, V., Larjo, A., Aho, T., Karp, M., Santala, V., 2011. Improved Triacylglycerol Production in Acinetobacter baylyi ADP1 by Metabolic Engineering. Microb Cell Fact 10, 36. 10.1186/1475-2859-10-36

Santala, S., Efimova, E., Santala, V., 2018. Dynamic decoupling of biomass and wax ester biosynthesis in Acinetobacter baylyi by an autonomously regulated switch. Metab Eng Commun 7, e00078. 10.1016/j.mec.2018.e00078

Santala, S., Santala, V., 2021. Acinetobacter baylyi ADP1—naturally competent for synthetic biology. Essays Biochem. 10.1042/EBC20200136

Scheller, H.V., Ulvskov, P., 2010. Hemicelluloses. Annu Rev Plant Biol 61, 263–289. 10.1146/annurev-arplant-042809-112315

Singh, K.D., Schmalisch, M.H., Stülke, J., Görke, B., 2008. Carbon Catabolite Repression in *Bacillus subtilis* : Quantitative Analysis of Repression Exerted by Different Carbon Sources. J Bacteriol 190, 7275–7284. 10.1128/JB.00848-08

Sokol, S., Millard, P., Portais, J.-C., 2012. influx_s: increasing numerical stability and precision for metabolic flux analysis in isotope labelling experiments. Bioinformatics 28, 687–693. 10.1093/bioinformatics/btr716

Stuani, L., Lechaplais, C., Salminen, A. V., Ségurens, B., Durot, M., Castelli, V., Pinet, A., Labadie, K., Cruveiller, S., Weissenbach, J., de Berardinis, V., Salanoubat, M., Perret, A., 2014. Novel metabolic features in Acinetobacter baylyi ADP1 revealed by a multiomics approach. Metabolomics 10, 1223– 1238. 10.1007/s11306-014-0662-x

Young, D.M., Parke, D., Ornston, L.N., 2005. OPPORTUNITIES FOR GENETIC INVESTIGATION AFFORDED BY ACINETOBACTER BAYLYI, A NUTRITIONALLY VERSATILE BACTERIAL SPECIES THAT IS HIGHLY COMPETENT FOR NATURAL TRANSFORMATION. Annu Rev Microbiol 59, 519–551. 10.1146/annurev.micro.59.051905.105823

Zhang, Y., Zhou, H., Tao, Y., Lin, B., 2020. Reconstitution of the Ornithine Cycle with Arginine:Glycine Amidinotransferase to Engineer *Escherichia coli* into an Efficient Whole-Cell Catalyst of Guanidinoacetate. ACS Synth Biol 9, 2066–2075. 10.1021/acssynbio.0c00138

Zhao, A., Hu, X., Wang, X., 2017. Metabolic engineering of Escherichia coli to produce gamma-aminobutyric acid using xylose. Appl Microbiol Biotechnol 101, 3587–3603. 10.1007/s00253-017-8162-3

Zhao, Z., Xian, M., Liu, M., Zhao, G., 2020. Biochemical routes for uptake and conversion of xylose by microorganisms. Biotechnol Biofuels 13, 21. 10.1186/s13068-020-1662-x

Zhou, H., Zhang, Y., Long, C.P., Xia, X., Xue, Y., Ma, Y., Antoniewicz, M.R., Tao, Y., Lin, B., 2024. A citric acid cycle-deficient Escherichia coli as an efficient chassis for aerobic fermentations. Nat Commun 15, 2372. 10.1038/s41467-024-46655-4

